# An affinity-matured human monoclonal antibody targeting fusion loop epitope of dengue virus with *in vivo* therapeutic potency

**DOI:** 10.1101/2020.10.03.324731

**Authors:** Tomohiro Kotaki, Takeshi Kurosu, Ariadna Grinyo, Edgar Davidson, Siti Churrotin, Tamaki Okabayashi, Orapim Puiprom, Kris Cahyo Mulyatno, Teguh Hari Sucipto, Benjamin J. Doranz, Ken-ichiro Ono, Soegeng Soegijanto, Masanori Kameoka

## Abstract

Dengue virus (DENV), from the genus *flavivirus* of the family *flaviviridae*, causes serious health problems globally. Human monoclonal antibodies (HuMAb) can be used to elucidate the mechanisms of neutralization and antibody-dependent enhancement (ADE) of DENV infections, leading to the development of a vaccine or therapeutic antibodies. Here, we generated eight HuMAb clones from an Indonesian patient infected with DENV. These HuMAbs exhibited the typical characteristics of weak neutralizing antibodies including high cross-reactivity with other flaviviruses and targeting of the fusion loop epitope (FLE). However, one of the HuMAbs, 3G9, exhibited strong neutralization ability (NT_50_ < 0.1 µg/ml) and possessed a high somatic hyper-mutation rate of the variable region, indicating affinity-maturation. Administration of this antibody significantly improved the survival rate of interferon-α/β/γ receptor knockout C57BL/6 mice after a lethal DENV challenge. Additionally, Fc-modified 3G9 molecules that had lost their in vitro ADE activity showed significantly enhanced therapeutic potency *in vivo* and competed strongly with an ADE-prone antibody *in vitro*. Taken together, the affinity-matured FLE-targeting antibody 3G9 exhibits several promising features for therapeutic application including a low NT_50_ value, potential for pan-flavivirus infection treatment, and suppression of ADE. This study demonstrates the therapeutic potency of affinity-matured FLE-targeting antibodies.

## Introduction

Many clinically-important mosquito-borne viruses including dengue virus (DENV), Japanese encephalitis virus (JEV), West Nile virus (WNV), and Zika virus (ZIKV) belong to the genus *flavivirus* of the family *flaviviridae* [1]. Of these, DENV causes the most serious health problems world-wide, in terms of the number of patients and fatalities. Infection with any of four serotypes of dengue virus (DENV-1 to DENV-4) causes dengue fever and dengue hemorrhagic fever [2], or dengue and severe dengue, as classified by the World Health Organization [3]. An estimated 390 million cases of DENV infection occur annually world-wide [4]. Of these, 100 million people develop symptomatic dengue and 21,000 die [4, 5].

One of the mechanisms hypothesized to cause a higher risk of severe dengue is the antibody dependent enhancement (ADE) of the infection, whereby pre-existing anti-DENV antibodies induced by the primary infection or vaccination facilitate subsequent DENV infections of Fc receptor-positive cells like macrophages [6]. Thus, DENV antibodies exhibit two conflicting activities: neutralization and ADE. Neutralization suppresses viremia, resulting in protection against DENV infection, while ADE increases viremia, and this is associated with severe dengue [7]. This phenomenon may increase the risk of developing severe dengue disease among seronegative people who have received vaccinations of CYD-TDV, which is the only licensed dengue vaccine [8, 9]. In addition, antibodies induced by ZIKV infection are reported to enhance DENV infections by ADE *in vitro*, and *vice versa* [10, 11]. ADE complicates dengue pathogenesis, and, thus, forms an obstacle to developing a fully effective dengue vaccine and prophylactic or therapeutic antibodies.

The DENV genome encodes three structural proteins (capsid [C], premembrane/membrane [prM/M], and envelope [E]) and seven non-structural proteins [1]. The viral particle is assembled in the lumen of the endoplasmic reticulum, where nucleocapsid (viral RNA complexed with the C protein) is incorporated into the lipid bilayer containing prM and E proteins [1]. As this immature viral particle traffics through the trans-Golgi network, a host serine protease (furin) cleaves the prM protein from the immature virus, resulting in maturation (the ability to infect). This maturation step occasionally remains incomplete, resulting in a mixture of virus particles at different states of maturity [12, 13]. Virus particles exhibit conformational dynamics referred to viral ‘breathing’ [14]. The maturity and breathing of virions have an impact on the recognition of antibodies and, thereby, affect their neutralizing and enhancing activities [15, 16].

The E protein is the major target of neutralizing antibodies, since it is located on the surface of a DENV virion [17]. Three domains (domains I, II, and III; DI, DII and DIII) have been identified in the E protein structure [18]. Each DENV particle contains 180 monomers of E protein that form 90 E-dimers [19].

The neutralization and ADE activities of antibodies are determined by the epitope of the virus [20, 21]. Antibodies targeting the fusion loop epitope (FLE) or bc loop on DII generally exhibit low levels of neutralization, high ADE, and high cross-reactivity to flaviviruses. These antibodies are extensively induced during secondary DENV infections, because the epitopes are highly conserved among the flaviviruses [22, 23]. Antibodies that bind to E-dimers, quaternary-structure epitopes, or the hinge regions of DI-DII exhibit strong neutralization ability by blocking viral conformational changes and membrane fusion [24–26]. Indeed, antibodies that recognize complex epitopes account for much of the virus neutralizing activity that occurs in the serum of convalescing patients [24,25,27]. However, many of these antibodies are serotype-specific. Antibodies that target domain III are serotype-specific and show higher neutralizing activity than those targeting domains I-II, although domain III-targeting antibodies are not predominantly produced in humans [25,27,28].

Human monoclonal antibodies (HuMAbs) could be useful tools for elucidating the mechanisms of neutralization and ADE, information that is required for vaccine development. In addition, HuMAbs can be used for prophylactic or therapeutic purposes. Several groups have been successful in generating HuMAbs against DENV, using various methods [24–26, 29]. Here, using newly developed SPYMEG cell technology, we generated eight anti-DENV HuMAb clones from an Indonesian patient with dengue [30, 31].

## Results

### Hybridoma preparation and details of HuMAbs

We established eight HuMAb clones from a blood specimen of an Indonesian patient with dengue using SPYMEG cells that belong to a human hybridoma fusion partner cell line [30, 31]. The patient was diagnosed with acute dengue fever (2 days after onset); the blood specimen was found to be anti-DENV IgM/IgG- and NS1-positive, indicating a secondary infection. The DENV serotype could not be determined by RT-PCR, using RNA extracted from the patient serum [32]. All HuMAbs were of the IgG1 subtype.

### Neutralization activity

All HuMAbs exhibited neutralizing activity against all four serotypes of DENV prototype strains and Indonesian isolates (Table 1, Fig.1A). DENV-2, in particular, was strongly neutralized; the NT_50_ was lower than 0.1 µg/ml. All of the HuMAbs also neutralized JEV. Overall, HuMAbs 1F11 and 3G9 showed stronger neutralizing activity. Thus, these two promising HuMAbs were subjected to neutralization tests using single-round infectious particles (SRIPs) containing prM-E proteins of ZIKV and WNV, due to the unavailability of the infectious viruses [33, 34]. Again, 1F11 and 3G9 neutralized ZIKV and WNV particles as efficiently as DENV (Fig. 1B). These data indicated that these HuMAbs were pan-flavivirus broadly-neutralizing antibodies.

**Figure 1.**
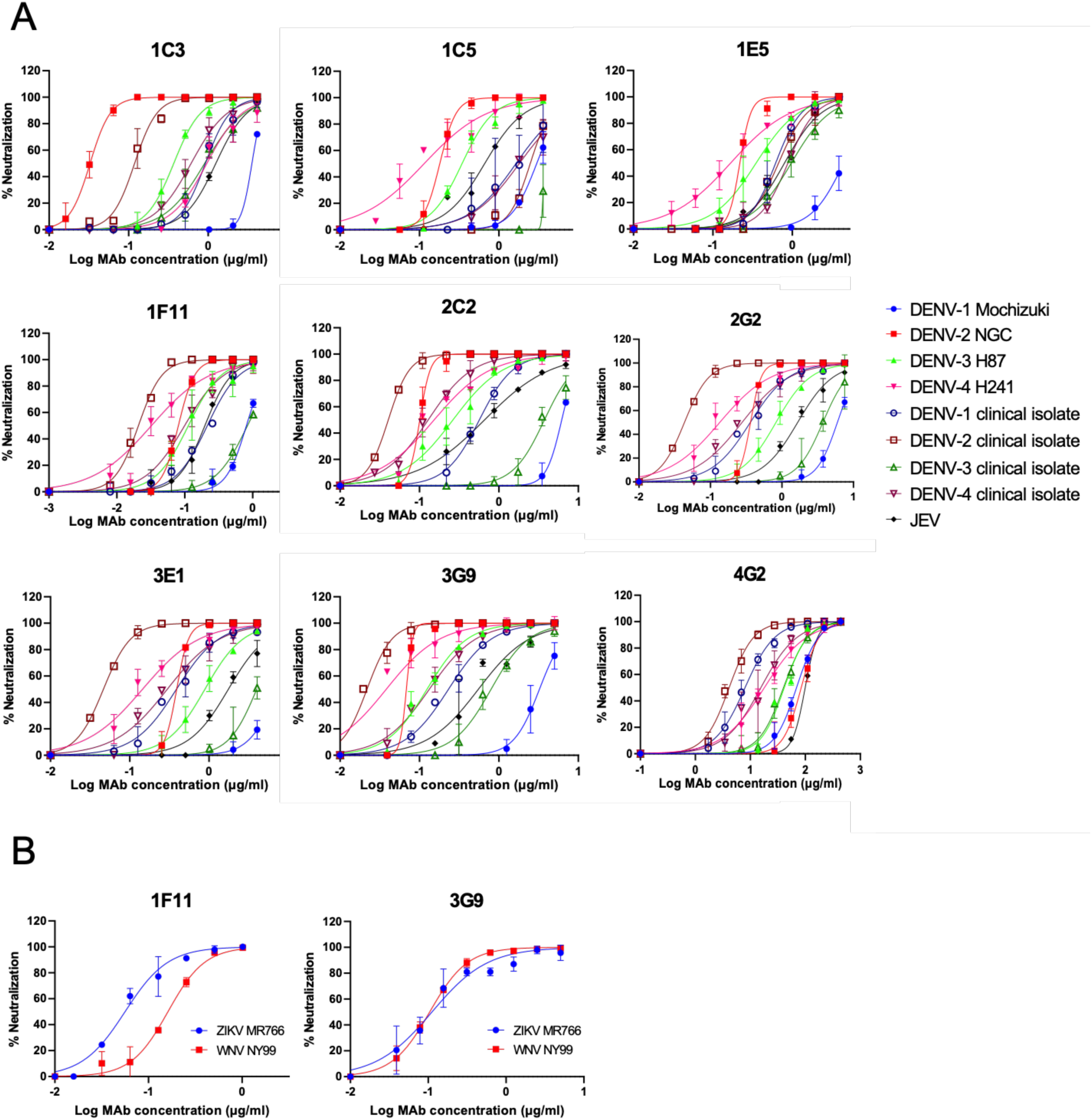
Neutralization of DENV, JEV, WNV, and ZIKV. A) Neutralizing activity against DENVs and JEV. B) Neutralizing activity against SRIPs containing prM-E of WNV and ZIKV. Average and SDs of two independent experiments are shown.

**Table 1.** NT_50_ values of the HuMAbs.

|  |  | Hybridoma clones |  |  |  |  |  |  |  |
| --- | --- | --- | --- | --- | --- | --- | --- | --- | --- |
|  |  | 1C3 | 1C5 | 1E5, | 1F11 | 2C2 | 2G2 | 3E1 | 3G9 |
| Indonesian isolates | DENV-1 | 0.90 | 1.89 | 0.65 | 0.24 | 0.59 | 0.21 | 1.00 | 0.24 |
|  | DENV-2 | 0.12 | 2.67 | 0.69 | 0.02 | 0.04 | 0.02 | 0.05 | 0.02 |
|  | DENV-3 | 0.87 | >3.60 | 0.92 | 1.29 | 3.39 | 1.99 | 2.41 | 0.73 |
|  | DENV-4 | 0.64 | 1.70 | 0.89 | 0.02 | 0.12 | 0.13 | 0.24 | 0.13 |
| Prototype strains | DENV-1 | 3.46 | 3.07 | >3.86 | 0.84 | 6.17 | 3.33 | 3.48 | 3.45 |
|  | DENV-2 | 0.03 | 0.18 | 0.21 | 0.09 | 0.10 | 0.20 | 0.11 | 0.06 |
|  | DENV-3 | 0.38 | 0.34 | 0.38 | 0.11 | 0.30 | 0.44 | 0.62 | 0.12 |
|  | DENV-4 | 0.89 | 0.09 | 0.17 | 0.03 | 0.11 | 0.06 | 0.22 | <0.04 |
| Other flaviviruses | JEV | 1.33 | 0.74 | 0.87 | 0.21 | 0.71 | 0.85 | 0.44 | 0.47 |
|  | WNV* | - | - | - | 0.17 | - | - | - | 0.11 |
|  | ZIKV* | - | - | - | 0.05 | - | - | - | 0.11 |

Average of two independent neutralization tests are shown (µg/ml).
\*The WNV and ZIKV results were determined using SRIPs containing the prM-E genes of each virus.

### Stage of HuMAb neutralization

Flavivirus infections can be neutralized by several mechanisms including the inhibition of receptor binding, inhibition of membrane fusion, and aggregation of virion particles [35]. A time of addition assay, using Vero cells and the DENV-2 New Guinea C (NGC) strain, was conducted to determine the neutralization mechanism used by the HuMAb clones [36]. The antibodies all neutralized DENV-2 during the pre-adsorption assay but not the post-adsorption assay (Table 2). This suggested that our HuMAbs blocked pre-adsorption steps including viral adsorption but not post-adsorption steps such as viral membrane fusion or conformation change.

**Table 2.** NT_50_ values from the preadsorption and postadsorption assays.

|  | Hybridoma clones |  |  |  |  |  |  |  |
| --- | --- | --- | --- | --- | --- | --- | --- | --- |
|  | 1C3 | 1C5 | 1E5 | 1F11 | 2C2 | 2G2 | 3E1 | 3G9 |
| Pre-adsorption | 0.04 | 0.30 | 0.47 | 0.06 | 0.16 | 0.43 | 0.23 | 0.07 |
| Post-adsorption | >8.3 | >7.2 | >0.95 | >2.0 | >13.9 | >8.1 | >15.1 | >5.9 |

Averages of two independent neutralization tests are shown (μg/ml).

### Epitope mapping

The epitope target of an antibody influences its neutralization potency and mechanism. Therefore, epitope mapping was conducted using HEK293 cells transfected with a DENV2 prM-E mutant library, which possess a single alanine substitution at each residue of prM-E (661 mutants in total) [37]. Four antibodies (1C3, 1F11, 3E1, and 3G9) were tested against all the mutants, while the other four antibodies (1C5, 1E5, 2C2, and 2G2) were tested only against selected mutants with an FLE mutation, because of the availability of materials. The data are shown in Supplementary Fig. S1. The critical residues that abolished the binding ability of the HuMAbs tested are presented in Fig. 2 and Table 3. The W101A mutation was critical to the activity of all eight HuMAbs. In addition to W101A, L107A and/or F108A were responsible for binding. All of the critical residues were located on the fusion loop of the E protein, which is highly conserved among the flaviviruses. These data substantiated the breadth of neutralizing ability exhibited by the HuMAbs considered in this study. Our antibodies were typical FLE antibodies, which are dominantly produced when a secondary dengue-associated infection occurs.

**Figure 2.**
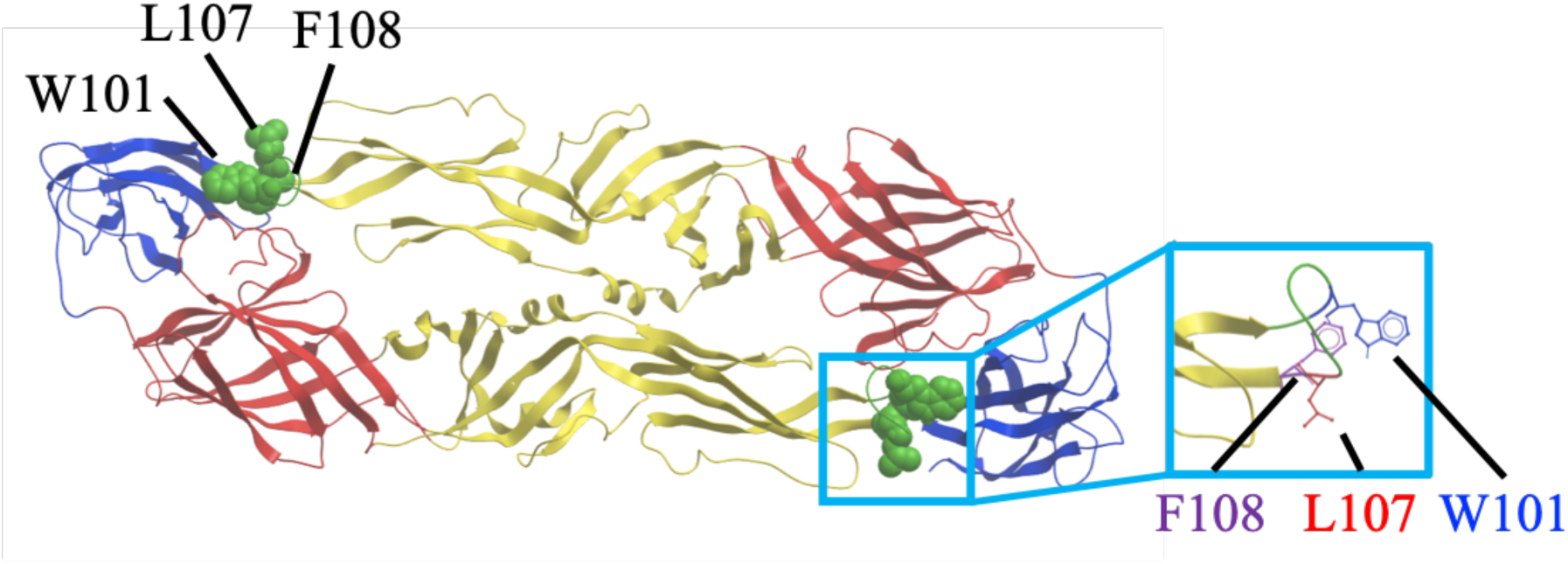
Deduced epitope locations on a ribbon diagram of the DENV-2 E protein. The deduced epitope locations were plotted on a DENV-2 E dimer, based on the data provided by the Protein Data Bank accession number 1OAN. DI, DII and DIII are indicated in red, yellow, and blue, respectively. The fusion loop region (98–110) is colored in green. Mutations affecting reactivity with the HuMAbs are shown as green spheres and in the magnified square.

**Table 3.** Critical mutations that abolished HuMAb binding.

|  | Hybridoma clones |  |  |  |  |  |  |  |
| --- | --- | --- | --- | --- | --- | --- | --- | --- |
|  | 1C3 | 1C5 | 1E5 | 1F11 | 2C2 | 2G2 | 3E1 | 3G9 |
| Critical<br>Residue | W101A | W101A | W101A | W101A | W101A | W101A | W101A | W101A |
|  | L107A | L107A | L107A | F108A | F108A | F108A | F108A | F108A |
|  |  | F108A |  |  |  |  |  |  |

The generation of escape mutants was attempted to further investigate the epitope, by passaging DENV-2 in the presence of the HuMAbs in Vero cells and then sequencing the E protein gene in any surviving viruses. However, we did not identify any mutations in the E protein of surviving viruses.

### ADE activity

Targeting of the FLE is typical of weak-neutralizing and high-ADE antibodies [22, 23]. We measured ADE activity using semi-adherent K562 cells [38]. Although the HuMAb clones exhibited strong neutralizing activity in Vero cells, strong ADE activity was observed in our assay system (Fig. 3). DENV-2 was neutralized at high concentrations of the HuMAb clones, while the other strains, DENV-1, −3, and −4, were not neutralized. These data suggested that the cloned HuMAbs could contribute to disease enhancing activity.

**Figure 3.**
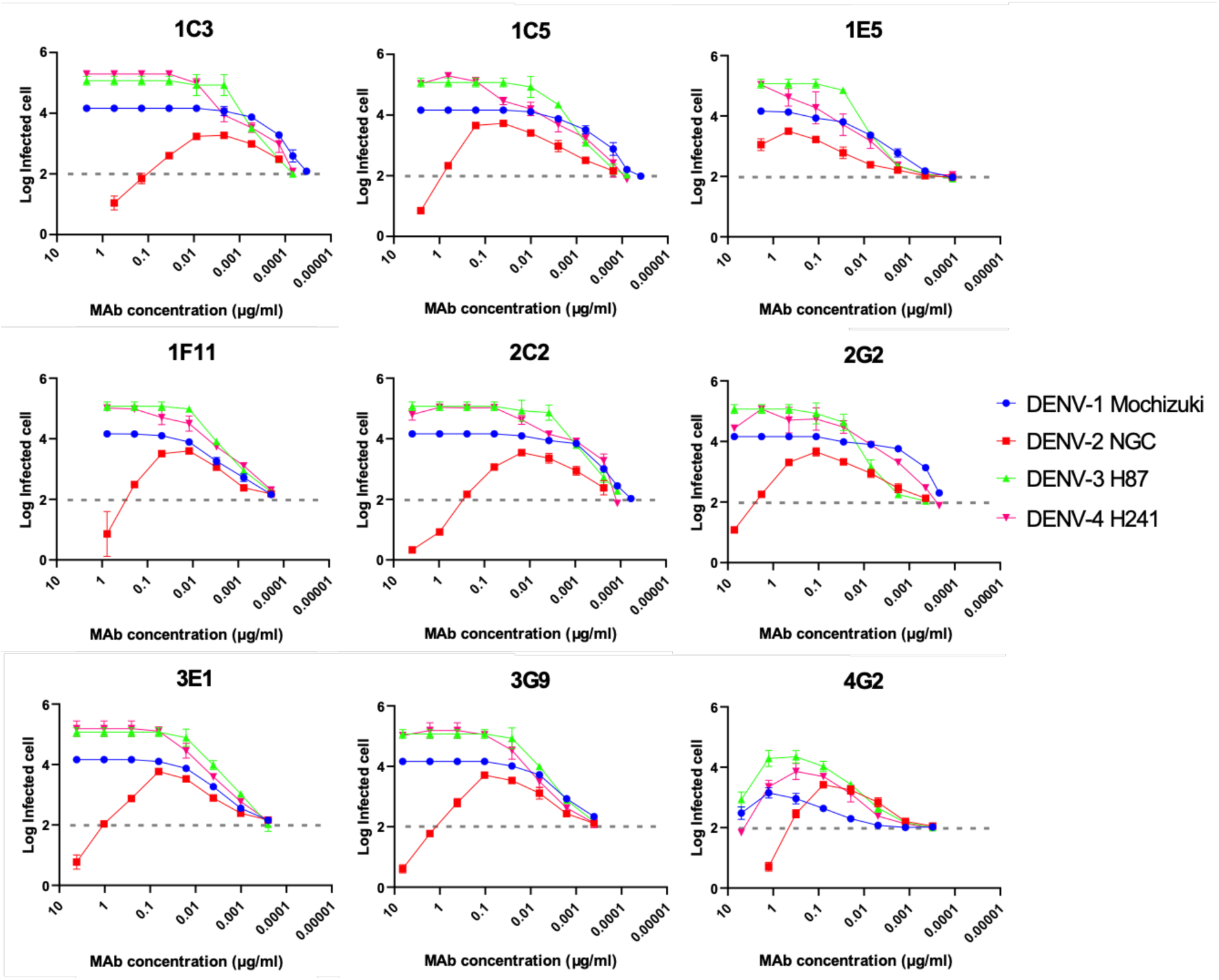
ADE activity of the HuMAbs. The DENV-2 NGC strain and K562 cells were used. Dotted lines indicate the baseline of the infected cells in the control (100 infected cells; 2.0). Each data point represents the mean obtained from two independent ADE assays with SDs.

### Recombinant IgG construction and evaluation

Although the cloned antibodies were potent in terms of low NT_50_ in Vero cells, high ADE was observed in K562 cells. However, these antibodies could potentially be used in therapeutic applications, by modifying the effector function and disrupting the ADE activity. Thus, one of three mutations [L234A + L235A (LALA), D265A, and N297A] was introduced to the Fc region of the best neutralizing HuMAb, 3G9, to disrupt the interaction with Fc receptors [39]. We chose to test three mutations, because the effect of Fc-modification during the *in vivo* animal experiment varied depending on the study group [40–45]. These three mutations are described as being the same in terms of the effect on loss of binding to Fc receptors and should not compromise antibody neutralizing potency [46] or shorten the half-life significantly [47]. However, N297A abolishes N-linked glycosylation at N-297, while the LALA and D265A mutants would be fully glycosylated. Additionally, LALA and N297A reduce complement component 1q (C1q) binding, whereas D265A does not compromise complement-dependent cytotoxicity (CDC), which is related to virus clearance [20, 39]. The difference in the mutations may cause subtle changes in the IgG phenotype, resulting in differences in *in viv*o efficacy [47].

3G9 was chosen to make recombinant antibodies, because it showed higher neutralizing activity and *in vivo* protection (see below). The recombinant Fc-modified 3G9 antibodies neutralized DENV-2 at comparable level to that of the 3G9-original in Vero cells (Table 4, Fig. 4A). As expected, the recombinant 3G9 antibodies did not show ADE activity in K562 cells, even at a sub-neutralizing concentration (Fig. 4B).

**Figure 4.**
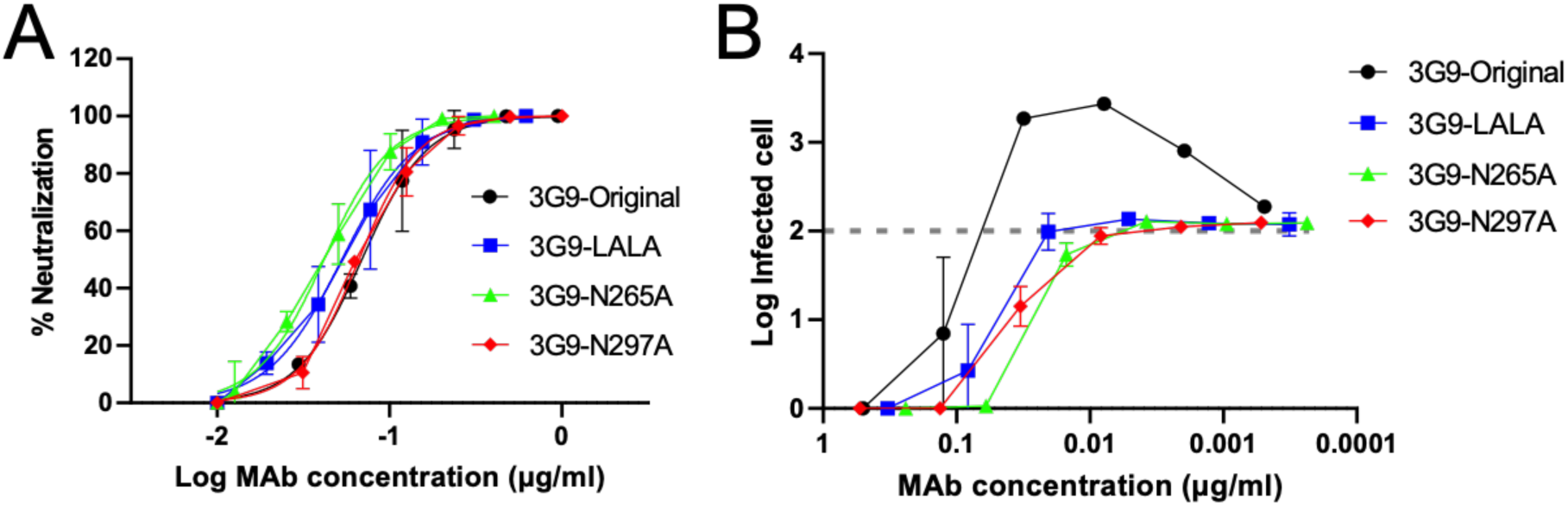
Neutralizing and ADE activities of the Fc-modified 3G9. A) Neutralizing activity against DENV-2 NGC strain using Vero cells. B) ADE activity against DENV-2 using K562 cells. Average and SDs of two independent experiments are shown.

**Table 4.** NT_50_ values of Fc-modified 3G9.

|  | 3G9-Original | 3G9-LALA | 3G9-N265A | 3G9-N297A |
| --- | --- | --- | --- | --- |
| NT<sub>50</sub> (µg/ml) | 0.069 | 0.053 | 0.042 | 0.066 |

Averages of two independent neutralization tests are shown (µg/ml).

### HuMAb protection in vivo

Because the neutralization and ADE activities *in vitro* did not always correlate with protection *in vivo*, we tested the ability of antibodies to protect against DENV infection in an animal model. We tested the ability of 3G9, 1F11 (another potent HuMAb), and Fc-modified 3G9 to protect interferon-α/β/γ receptor knockout (IFN-α/β/γR KO) C57BL/6 mice challenged with DENV-3 [48]. DENV-3 was chosen because of the availability of the lethal infection mouse model, in which DENV infection causes vascular leakage without showing neurologic disorder [48]. Antibodies were injected intraperitoneally (i.p.) after one day of virus challenge. Mice administered with an isotype control IgG died within 4 days postinfection (Fig. 5). 1F11 exhibited almost no protection, while the original unmodified 3G9 significantly prolonged survival, although all mice died within 20 days post-infection (*p* < 0.01). All three recombinant 3G9 antibodies significantly enhanced the survival rate when compared with the original 3G9 (*p* < 0.05). There were no significant differences among the three recombinant antibodies (*p* > 0.05). These results indicated that the reduced ADE shown by these modified antibodies promotes their effectiveness *in vivo* and suggests that recombinant 3G9 antibodies are promising therapeutic agents.

**Figure 5.**
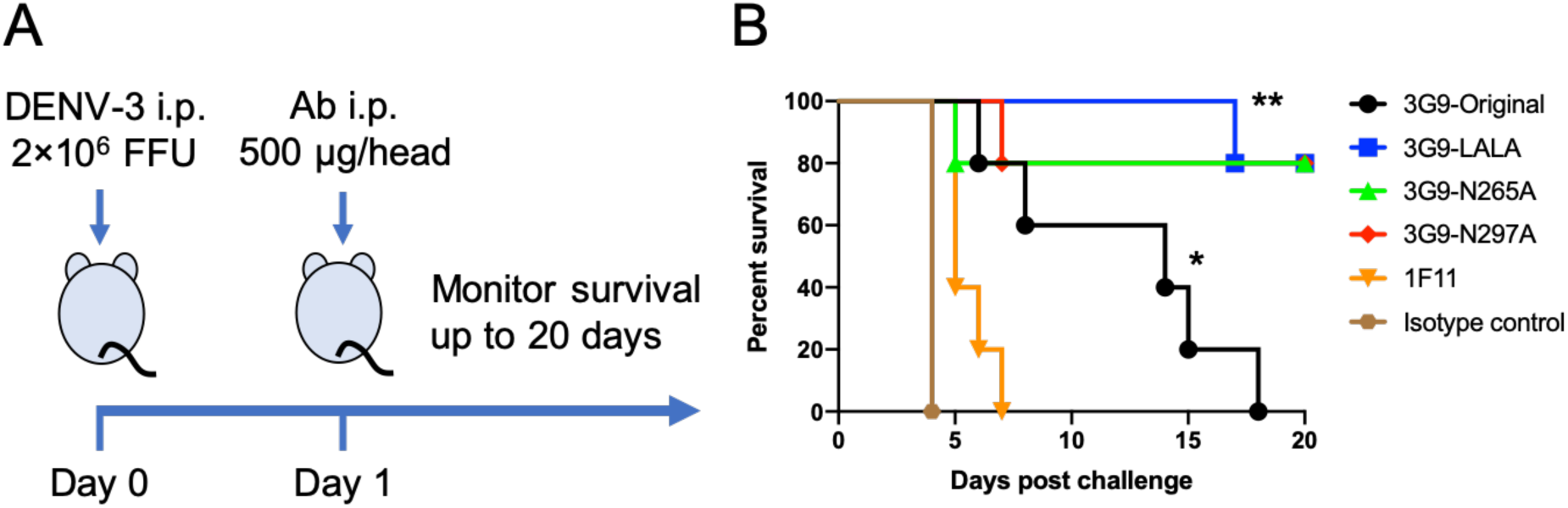
*In vivo* efficacy test. A) Scheme of the animal challenge experiment. B) Survival of infected mice after viral challenge. A group of five mice were infected with 2.0 × 10^6^ FFU of DENV-3. Statistical significance was analyzed using a Kaplan-Meier method. \**P <* 0.01 for the comparison between isotype control and 3G9-original. \*\**P <* 0.05 for the comparison between 3G9-original and the Fc-modified 3G9. There was no significant difference among the Fc-modified 3G9 (*P >* 0.05).

### Competition assay using ELISA and ADE assays

Our animal experiment was carried out without ADE in the post-infection treatment model. Thus, it remained unclear whether the Fc-modified 3G9 could suppress ADE *in vivo*. An antibody highly competitive to 4G2 (low avidity FLE antibody) has been reported to protect mice from lethal DENV infection with ADE [49]. Since we had not established an *in vivo* ADE infection model, competition assays were conducted. Competitive ELISA showed that 3G9 strongly competed with 4G2 but not with other mouse monoclonal antibodies (7F4 and 15C12, targeting the central part of domain II and the A strand of domain III, respectively) (Fig. 6A) [20]. 3G9 inhibited 4G2 binding by more than 50% when both were applied to the assay at the same concentration.

**Figure 6.**
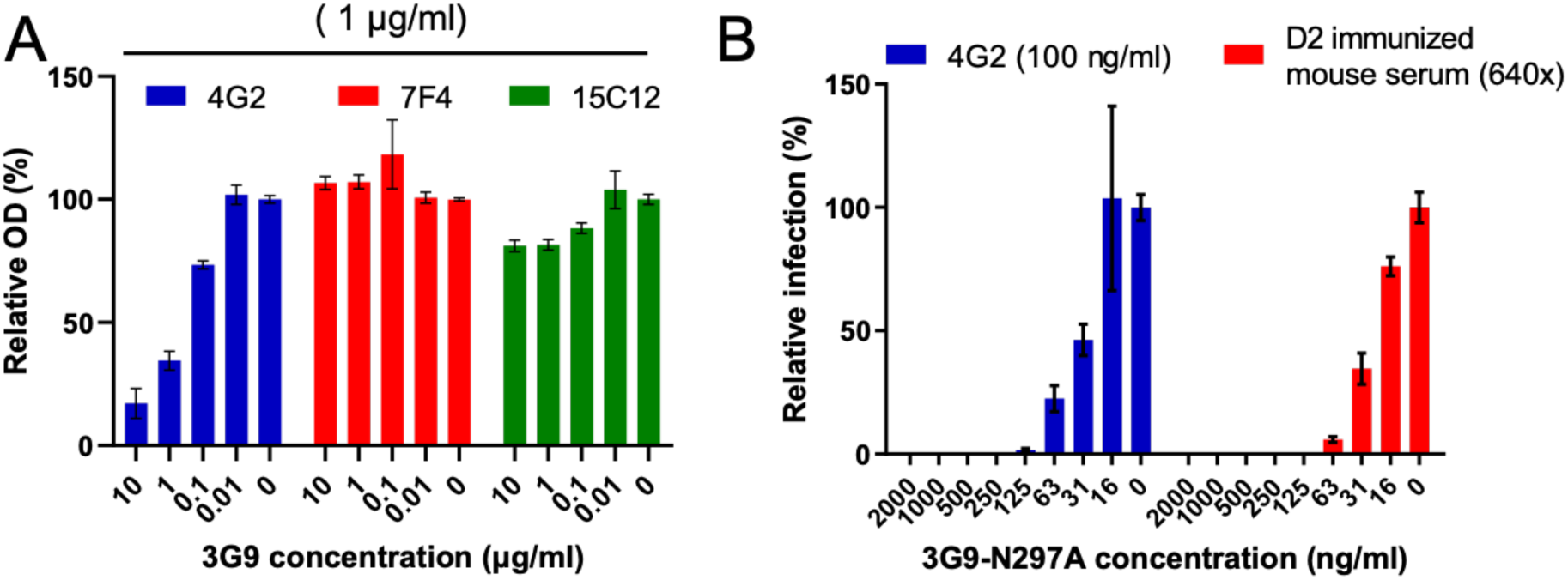
Competition ELISA and ADE assay. A) Competition ELISA. Mouse monoclonal antibodies 4G2, 7F4, or 15C12 at 1 µg/ml were mixed with serially diluted 3G9 and incubated in a DENV-coated ELISA plate. The OD relative to that of the no competition well (without 3G9) is shown with SDs of triplicate experiments. B) Competition ADE. 100 ng/ml of 4G2 and 1:640 diluted mouse serum, which showed the peak level of enhancement, were mixed with DENV-2 NGC and serially diluted 3G9-N297A. The number of infected cells relative to that of the no competition wells is shown with SDs of triplicate experiments.

It has been reported that the suppression of ADE caused by pre-existing antibodies *in vitro* could be an indicator of *in vivo* protection efficacy [49]. Thus, a competitive ADE assay was performed using 4G2 and DENV2-immunized mouse serum, which showed peak levels of ADE at 100 ng/ml and a 1:640 dilution, respectively (Fig. 3, Fig. S2). These dilutions were used, therefore, for the subsequent competitive ADE assay. 3G9-N297A suppressed the peak ADE caused by 4G2 and D2-immunized mouse serum by 50% at only 30 ng/ml (Fig. 6B). A previous study indicated that an antibody that strongly reduced ADE infection (>50%) at 1000 ng/ml offered good therapeutic efficacy in an *in vivo* ADE model [49]. The competition intensities found in the current study were higher than the threshold for *in vivo* protection quoted in the previous report, although direct comparison is impossible.

### Immunogenetic analysis

Even though 3G9 is an FLE antibody, it showed high potency in terms of NT_50_ values and *in vivo* protection. Therefore, we analyzed the sequences of the 3G9 VH and VL regions using the IMGT tool to identify the closest VH and Vλ germline genes. The results indicated that the VH gene was derived from IGHV3-23*02 and the Vλ gene from IGLV7-46*01 (Table 5). Somatic hyper-mutation (SHM) rates of the VH and Vλ genes were 14.3% and 7.1%, respectively. The high SHM rate of the VH regions indicated the levels of affinity-maturation during the secondary infection [42].

**Table 5.**
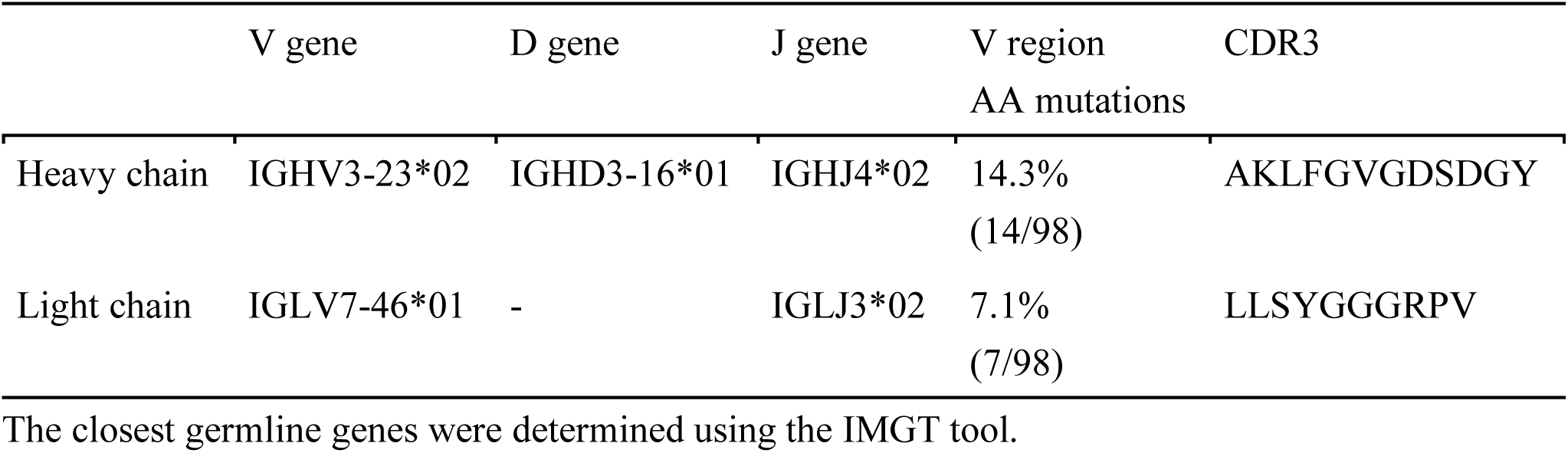
Immunogenetic analysis of 3G9.

## Discussion

To date, no specific therapeutic agent against flavivirus infection has been made available. Considering the current success of antibody therapy against respiratory syncytial virus and Ebola virus infections, antibody therapy for dengue is a promising target. DENV-neutralizing HuMAbs have been reported using various techniques and patient backgrounds [24-26,29,31]. The objective of this study was to establish therapeutic antibody candidates against dengue using the newly developed SPYMEG cell technology [30, 31]. Multiple experiments were attempted using blood samples from dengue patients with various backgrounds. We successfully established eight HuMAb clones from a single dengue patient. The patient whose hybridoma was successfully established was in the acute phase of secondary dengue infection, indicating that the B cells were highly activated and IgG repertoire was abundant. The SPYMEG cell fusion method was highly successful when the patient was in the acute infection phase, which was consistent the results of a previous study [31]. The circulating serotypes and/or genotypes of DENV vary depending on the region and year [50, 51]. These antibodies, therefore, could be a potent therapeutic agent for use against DENV strains circulating in Southeast Asia including Indonesia, which is one of the largest dengue-endemic countries.

Several groups have found potent neutralizing HuMAbs against DENV. In general, highly potent neutralizing antibodies are serotype-specific, bind to E-dimers or quaternary-structure epitopes, and inhibit both pre- and post-attachment steps [52]. In contrast, weak neutralizing antibodies are highly cross-reactive, bind to E-monomer FLEs, and inhibit the only pre-attachment step. All of the HuMAbs cloned in this study exhibited characteristics of weak neutralizing antibodies. However, they showed considerably low NT_50_ values, especially against DENV-2. The NT_50_ values were <0.1 µg/ml, as low as those of previously reported highly potent neutralizing antibodies [52]. However, these values cannot be compared directly, because the neutralizing antibody titer is influenced by the host cell type, degree of viral maturation, virion breathing, and other assay conditions [53, 54]. Thus, 4G2 (a low neutralizing FLE antibody) was used to standardize the NT_50_ assay. One of the HuMAb clones, 3G9, exhibited a NT_50_ that was 1000-fold lower than that of 4G2 against DENV-2 (Table 1). This was comparable to the findings of a previous study on potent neutralizing antibodies [55], corroborating the suggestion that 3G9 is a promising therapeutic antibody.

High neutralizing potency, especially against DENV-2, is beneficial, because the only licensed dengue vaccine (CYD-TDV) confers low-level protection against DENV-2 [56, 57]. Cross-reactive neutralization is reported to play an important role in DENV-2 protection, while serotype-specific neutralization is key to protection against other serotypes [58]. The HuMAbs described in this study could compensate for the drawbacks associated with the current dengue vaccine.

3G9 also neutralized other flaviviruses, with NT_50_ values of around 0.1 µg/ml. As ZIKV is less susceptible to FLE antibodies, due to high thermostability and less virion breathing [59, 60], a potent neutralizing antibody targeting both DENV and ZIKV could be promising, considering their co-circulation in the environment and the occurrence of ADE in both diseases [10, 11]. The HuMAbs identified in this study may be applied therapeutically in the treatment of pan-flavivirus infections.

All of the HuMAbs investigated were sensitive to W101A mutation, which is a typical characteristic of FLE antibodies [26]. Although all of the HuMAbs recognized the fusion loop, the epitope type was further classified into three groups (Table 3). Group 1 (1C3 and 1E5) recognized W101 and L107; group 2 (1C5) recognized W101, L107, and F108; group 3 (1F11, 2C2, 2G2, 3E1, and 3G9) recognized W101 and F108. Group 1 and 2 antibodies, which recognize the L107 residue, were 4–50 times less potent than those of group 3. Accordingly, L107 could be a marker for low neutralization activity, and F108 might be critical to viral fusion or internalization.

Other groups have reported potent neutralizing antibodies that target FLEs. For example, 2A10G6, targeting D98, R99, and W101 motifs within the fusion loop, has a broad neutralizing capability against all four serotypes of DENV and protects mice against lethal challenges from DENV and WNV [61]. 753C12, targeting both the fusion loop and adjacent bc-loop, exhibits low NT_50_ values [62]. Furthermore, E60 is a potent neutralizing FLE antibody [49, 63]. These reports substantiate the potential of FLE antibodies.

No viral escape mutant was obtained, even after 5 viral passages in the presence of the HuMAbs in Vero cells. The fusion loop is highly conserved among flavivirus, and an escape mutation at this position would be lethal to the virus. This feature also could be an advantage to the development of therapeutics.

Generally, cross-reactive non-conformational epitopes are less neutralizing, while serotype-specific conformational epitopes are highly potent [52]. Most serotype-specific antibodies are induced at the primary infection stage, while cross-reactive FLE antibodies are induced exclusively at the secondary infection phase [22, 23]. Tsai et al reported that FLE antibodies derived from patients with a secondary infection are more potent than that those from patients with a primary infection [62]. Considering the high SHM rate (14.2%) on the V gene (Table 5), 3G9 was considerably affinity-matured, and, therefore, potent [42]. In addition, as a third or fourth DENV infection is less possible, affinity-matured FLE antibodies play an important role in viral protection [64].

Although the HuMAbs investigated in this study were promising in intensity and breadth of neutralization, strong ADE was observed. DENV-2 was slightly neutralized at a high concentration of IgG in the ADE assay, coinciding with the high level of neutralization activity in Vero cells. Meanwhile, the other serotypes of DENV were not neutralized. The ADE assay measures the balance of neutralizing and enhancing activities, and, thus, no neutralizing activity was observed [38]. The HuMAb clones were subtyped as IgG1, which show relatively higher levels of ADE activity [65]. These data indicate that modification of the Fc region is mandatory prior to therapeutic application.

Fc-modified 3G9 neutralized DENV at a similar level to the original antibodies, but ADE was not observed, even at sub-neutralizing levels *in vitro*. In addition, these recombinant antibodies significantly prolonged the survival of the infected mice, compared with the original antibody. Some groups have reported that disrupting the Fc-interaction significantly enhances survival rates [42,44,49], while other groups have demonstrated the attenuation of protection or no significant difference [41, 43]. Our study demonstrated a significantly improved survival rate and no difference among the three recombinant antibodies. These data suggested that disrupting Fc-interactions is promising, at least in mice, regardless of the mutations. We used IFN-α/β/γR knock-out C57BL/6 mice, which are genetically distinct from AG129 mice; the latter are commonly used for dengue virus challenge experiments. This difference did not appear to affect the phenotype of the mice infected with DENV [48]. However, it is still possible that the genetic background and viral strain may have resulted in a different outcome from the virus challenge experiment.

3G9 did not completely prevent death by viral infection in our model. This may reflect the fact that 3G9 showed less neutralizing activity to DENV-3 than the other DENV serotypes (Table 1). In addition, we challenged each mouse with a high amount of virus (2 × 10^6^ focus forming unit [FFU]). Nevertheless, Fc-modified 3G9 prevented the death of 80% of the mice at 20 days post-infection. This antibody would be expected to protect the mice against DENV-2 efficiently, considering the 2–4-fold lower NT_50_ value of DENV-2 (Table 1). Again, the results indicated that Fc-modified 3G9 is a good therapeutic candidate.

While our data suggests the importance of eliminating the ADE function of protective antibodies, a limitation of our study was the lack of animal experiments using an ADE model. Williams et. al. reported that the assay of antibody competition or ability to displace low-avidity FLE antibodies and reduce ADE *in vitro* could enable the prediction of *in vivo* therapeutic efficacy with ADE [49]. Our ADE competition assay data revealed that 3G9 competes with 4G2 and polyclonal immunized-mouse serum (Fig. 6). The observed degree of competitiveness was stronger than that shown previously by E60, a potent neutralizing FLE antibody [49], although direct comparison was impossible. An affinity-matured FLE antibody would be superior to other potent neutralizing antibodies targeting the DI-DII hinge region or DIII, in terms of competing with pre-existing ADE-prone FLE antibodies, which may warrant the suppression of ADE *in viv*o [49].

In conclusion, we have described a potent HuMAb, 3G9, which targets FLE. Fc-modified 3G9 displayed neutralizing potency *in vivo*. The affinity-matured FLE antibody has several features that make it appropriate for therapeutic application including a low NT_50_ value, potential for pan-flavivirus infection treatment, competition with pre-existing ADE-prone antibodies, and less potential for viral escape. This study reconfirms the therapeutic potency of affinity-matured FLE antibodies.

## Methods

### Ethical statement and patient recruitment

Blood samples were collected from patients with dengue at a private hospital in Surabaya, Indonesia. Signed informed consent was acquired from the patients or their parents upon collection of blood samples. This study was approved by the Ethics Committees of Airlangga University (Ethics Committee Approval Number: 24-934/UN3.14/PPd/2013) and Kobe University Graduate School of Medicine (Ethics Committee Approval Number: 784). Diagnosis data were obtained from medical records.

All animal experiments were performed at the National Institute of Infectious Diseases in Japan (NIID), in accordance with the guidelines for the care and use of laboratory animals at the NIID. The study was approved by the Animal Experiment Committee of the NIID (Ethics Committee Approval Number: 115064). Trained laboratory personnel performed the anesthesia of mice via i.p. injection of a mixture of medetomidine, midazolam, and butorphanol during viral inoculation and euthanasia by cervical dislocation.

### Hybridoma preparation

Approximately 5 ml of blood were obtained from one patient and the peripheral blood mononuclear cells were isolated by centrifugation using a Ficoll-Paque PLUS (GE Healthcare, Uppsala, Sweden). These cells were then fused with SPYMEG cells at a ratio of 10:1, to establish hybridomas, as described previously [30, 31].

### Cell lines

The SPYMEG cells and established hybridomas were maintained in Dulbecco’s modified Eagle medium (DMEM) supplemented with 15% fetal bovine serum (FBS) and 3% BM-condimed (Sigma Aldrich, St. Louis, MO) [18]. Vero cells were cultured in Eagle’s minimum essential medium (MEM) supplemented with 10% FBS and 60 µg/ml kanamycin. C6/36 cells were cultured in MEM supplemented with 10% FBS, nonessential amino acids, and 60 µg/ml kanamycin. HEK-293T cells were maintained in DMEM supplemented with 10% FBS. K562 erythroleukemia cells were cultivated in RPMI 1640 medium supplemented with 10% FBS, 100 units/ml penicillin, and 100 µg/ml streptomycin. FreeStyle 293-F cells (Invitrogen, Gaithersburg, MD) were cultured in FreeStyle 293 Expression Medium (Thermo Fisher Scientific, Waltham, MA).

### Viruses

DENV-1 genotype I, DENV-2 cosmopolitan genotype, DENV-3 genotype I, and DENV-4 genotype II, which are typical isolates found in Surabaya, Indonesia, were used for the antibody characterization [50, 51]. Sequence information is available upon request. In addition, the Mochizuki strain of DENV-1, NGC strain of DENV-2, H87 strain of DENV-3, H241 strain of DENV-4, and Nakayama strain of JEV were used as prototype viruses [66]. The DENV-3 strain P12/08, derived from patients infected with DENV-3 in Thailand in 2008, was used in the animal experiments [48]. Viruses were propagated in C6/36 cells and stored at - 80°C until use.

### Hybridoma screening by ELISA

Rabbit serum immunized with the DENV-2 NGC strain was coated onto Nunc Maxisorp ELISA plates (Thermo Fisher Scientific, Waltham, MA). Then, the DENV-2 Indonesian strain (inactivated with Tween 20), hybridoma culture (or DMEM as a negative control), alkaline phosphatase (AP) conjugated anti-human IgG (Abcam, Canbridge, UK), and p-nitrophenyl phosphate (PNPP) (Nacalaitesque, Kyoto, Japan) were serially incubated, and the absorbance was measured at 415 nm. The wells showing higher values than the average + 3SD of the negative controls were considered to be positive.

### Isotyping and quantification of HuMAbs

HuMAbs were isotyped using anti-human IgG1, IgG2, IgG3, or IgG4 by ELISA (Abcam, Canbridge, UK). HuMAbs were coated onto the ELISA plates. Then, murine anti-human IgG (anti-IgG1, IgG2, IgG3, or IgG4), AP-conjugated anti-mouse IgG, and PNPP were serially incubated, and the absorbance was measured at 415 nm. The subclass of each HuMAb was determined following the targeted identification of the first antibody’s subclass, which showed the highest value among IgG1 to IgG4. Human IgG was quantified using a Human IgG Quantification kit (RD Biotech, Besançon, France).

### Preparation of single round infectious particles

pCMV-JErep-fullC, a pcDNA3 plasmid containing JEV genes encoding the whole C and all the NS proteins, was transfected into 293T cells with a prM-E expression plasmid of the WNV NY99 strain or ZIKV MR776 strain, to prepare SRIPs [33, 34]. The culture media containing the SRIPs were harvested 3 days post-transfection. The harvested SRIPs were subjected to neutralization tests.

### Antibodies and immunized serum

D1-4G2 (anti-E protein, cross-reactive to the flavivirus group; American Type Culture Collection, Manassas, VA) and JE-2D5 (anti-JEV-NS1 protein) were used to detect virus-infected and SRIP-infected cells, respectively (see below) [33]. Two mouse monoclonal antibodies, 7F4 (targeting the central part of DII) and 15C12 (targeting the A strand of DIII), were used for the competition ELISA [20].

In addition, a mouse polyclonal antibody against DENV-2 (immunized mouse serum) was used for the competition ADE assay. Six-week old BALB/c mice were immunized three times with 100 µg of DNA vaccine (prM-E protein expression plasmid) intratibially, at two-week intervals [66]. Blood samples were collected one week after the third immunization.

### Titration of viral infectivity and neutralization test

Infective titers were determined in Vero cells on a 96-well plate, by counting the infectious foci after immunostaining (see below) and expressed as FFU.

Neutralizing tests were performed as described previously [66]. Briefly, flat-bottom 96-well plates were seeded with Vero cells (2 × 10^4^ cells/well). The following day, 100 FFU of virus and serially diluted antibody were mixed and incubated for 1 h at 37°C, followed by inoculation into the Vero cells. At 24 h post-infection, the cells were fixed and immunostained (see below). The neutralizing antibody titer was expressed as the minimum IgG concentration yielding a 50% reduction in focus number (NT_50_).

### Immunostaining

Immunostaining was performed as described previously [66]. Briefly, infected cells were fixed with acetone-methanol (1:1). These cells were incubated serially with the antibodies described above, biotinylated anti-mouse or -human IgG, a VECTASTAIN Elite ABC kit (Vector Laboratories, Burlingame, CA), and a VIP peroxidase substrate kit (Vector Laboratories, Burlingame, CA).

### Time of addition assay

A time of addition assay was carried out, as described previously [36]. For the pre-adsorption assay, approximately 100 FFU of virus were pre-incubated with serially diluted HuMAbs for 1 h at 4°C and then inoculated onto 2 × 10^4^ Vero cells on a 96-well plate. Then, the unadsorbed viruses and excess antibodies were washed out with PBS. The cells were then incubated for 24 h at 37°C, followed by immunostaining and focus counting.

For the post-adsorption assay, virus was added directly to the cells for 1 h at 4°C. Then, unadsorbed virus was removed by washing the cells with PBS three times, and bound virus was incubated with serially diluted HuMAbs for an additional hour at 4°C. The cells were then incubated for 24 h at 37°C, followed by immunostaining and focus counting. The results are expressed in the same way as for the neutralization assay.

### Epitope mapping

Epitope mapping was conducted as reported previously [37]. A DENV2 (strain 16681) prM-E expression construct was subjected to high-throughput ‘Shotgun Mutagenesis’, generating a comprehensive mutation library. Each prM-E residue of the construct was changed individually to alanine (alanine residues to serine). In total, 661 DENV2 mutants were generated (100% coverage of prM-E). HEK-293T cells were transfected with an expression vector for DENV-2 prM-E or its mutants, fixed with 4% paraformaldehyde, and intracellular MAb binding was detected using a high-throughput immunofluorescence flow cytometry assay. Antibody reactivity against each mutant protein clone was calculated relative to the reactivity of the wild-type protein. Each raw data point was background-subtracted and normalized to the value for reactivity with wild-type DENV2 prM-E. Mutations within clones were identified as critical to the MAb epitope if they did not support reactivity of the MAb (<20% of the MAb’s reactivity with wild-type prM-E) but did support reactivity of other conformation-dependent MAbs (>70% of reactivity with wild-type).

### Generation of escape mutants

The DENV-2 NGC strain was passaged 5 times in the presence of the HuMAbs in Vero cells. The concentration of HuMAbs was increased with passages, starting from NT_50_ to 5 times NT_50_. Surviving viruses were sequenced in the E region.

### ADE assay

ADE activity was measured using semi-adherent K562 cells and expressed as the number of infected cells [38]. Briefly, serial four-fold dilutions of antibody samples were incubated with 100 FFU virus for 2 h at 37°C in 96 well poly-L-lysine coating plates. The mixture was mixed with 1 × 10^5^ K562 cells and incubated for a further 2 days. After immunostaining, viral foci were counted manually. The baseline of the infected cells (without antibody) was 2.0 (100 FFU). The infected cell number fell lower than 2.0 when the virus was neutralized but rose higher when ADE occurred.

### Generation of Fc-modified recombinant antibodies

RNA was extracted from 5 × 10^6^–10 × 10^6^ hybridoma cells using TRIzol reagent (Invitrogen, Gaithersburg, MD). Heavy (H)- and light (L)-chain cDNAs containing the gene encoding the antibody-binding (Fab) region of 3G9 were amplified and sequenced, as previously reported [67]. Then, primer sets were designed to clone the Fab regions of H and L chains into pFUSE-hIgG1-Fc1 and pFUSE2-CLIg-hL2, respectively (InvivoGen, San Diego, CA). Gene cloning was performed following the manufacturer’s instructions (primer information is available upon request).

Then, three kinds of mutation [L234A/L235A(LALA), D265A, and N297A] that abolished Fc-Fc receptor interaction were introduced into the Fc region using a site-directed mutagenesis kit, following the manufacturer’s protocol (TOYOBO, Osaka, Japan) [39].

The plasmids containing H- or L-chain genes (50 µg each) were transduced to 1 × 10^8^ 293F cells using 293fectin reagent (ThermoFisher Scientific, Waltham, MA). After incubating the cells at 37°C for 4 days, the culture fluids were harvested and purified by protein G (GE Healthcare, Chicago, IL). The concentration of purified IgG was calculated by measuring the absorbance at 280 nm. Purified IgG was then used in neuralization tests, ADE assays, and animal experiments (see below).

### Animal experiments

Five or six 6-week old IFN-α/β/γR KO C57BL/6 mice per group were challenged i.p. with 2 ×10^6^ FFU of DENV-3 (P12/08) under anesthesia. Twenty hours post challenge, HuMAbs 1F11, 3G9, 3G9-LALA, 3G9-N265A, or 3G9-N297A (500 µg/mouse) were injected i.p., and the mice were observed for 20 days. Mice were euthanized for humane purposes if they showed apparent symptoms.

### Competition assays

Competition ELISA: The DENV-2 NGC strain was coated on ELISA plates, as described above. Then, 1 µg/mL of each type of mouse monoclonal antibody (4G2, 7F4, or 15C12) was mixed with serially diluted 3G9 (four 10-fold dilutions starting at 10 µg/mL) in a separate 96-well plate, and 100 µl of the mixture were added to each ELISA plate. The plates were then incubated with AP-conjugated anti-mouse IgG, followed by color development with PNPP. The relative optical density (OD) was expressed as the average OD of each sample divided by the OD of the non-competition control well (without 3G9). Competition ADE: Serially diluted 3G9-N297A (eight 2-fold dilutions starting at 2000 ng/ml) solutions were prepared in a separate 96-well plate, followed by the addition of 4G2 or DENV-2 immunized to the concentration that showed peak enhancement of DENV-2 infection in K562 cells. Then, 36 µl of the mixture were used for the ADE assay, as described above [38]. Relative infection was expressed as the average number of infected cells in each sample divided by the number of infected cells in the non-competition control well (without 3G9-N297A, around 1000 cells).

### Statistics

All error bars indicate standard deviations. All calculations were performed using GraphPad Prism 8 (GraphPad Software Inc.).

## Acknowledgements

This work was supported by a program from the Japan Initiative for Global Research Network on Infectious Diseases (J-GRID) through the Ministry of Education, Culture, Sport, Science and Technology in Japan, and the Japan Agency for Medical Research and Development (AMED); the Institute of Tropical Disease (ITD) and the Center of Excellence (COE) program from the Ministry of Research and Technology (RISTEK), Indonesia. pCMV-JErep-fullC and a plasmid containing prM-E gene of ZIKV were provided by Dr. Ryosuke Suzuki, NIID and Dr. Eiji Konishi, Osaka university, respectively. The funders had no role in study design, data collection and analysis, decision to publish, or preparation of the manuscript. The manuscript was proofread by Enago.

## Author contributions

T. Kotaki and M.K. conceived the study. T. Kurosu performed the animal experiments. A.G., E.D., and B.J.D performed the epitope mapping. S.C., K.C.M., T.H.S., and S.S. helped to obtain clinical specimens. S.C., T.O., and O.P. helped to generate the HuMAbs. K.-I. O. provided critical reagents (SPYMEG cells). T. Kotaki performed the other experiments and took the lead in writing the manuscript. T.Kurosu, E.D., T.O., and M.K. provided feedback and helped shape the research and manuscript.

## Additional information

### Data availability

The datasets generated during and/or analyzed during the current study are available from the corresponding author on reasonable request.

