## Supplementary information for "An affinity-matured human monoclonal antibody targeting fusion loop epitope of dengue virus with *in vivo* therapeutic potency"

A

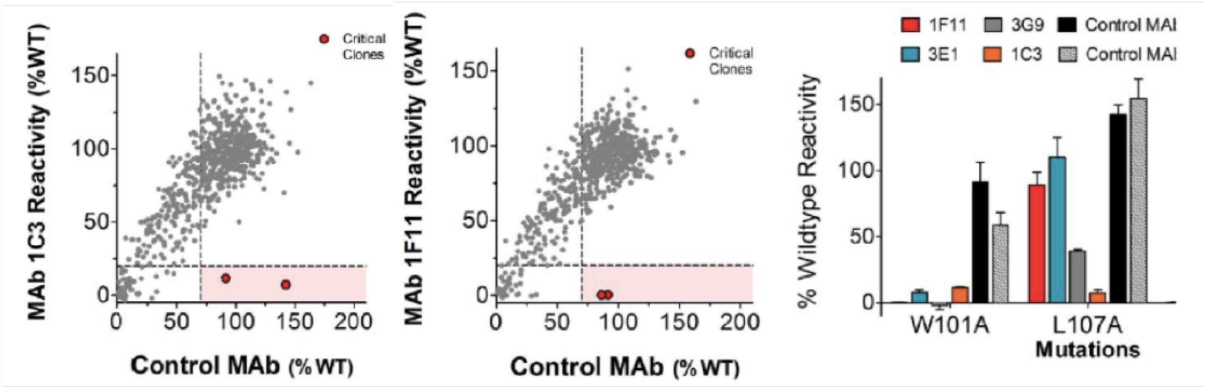


B

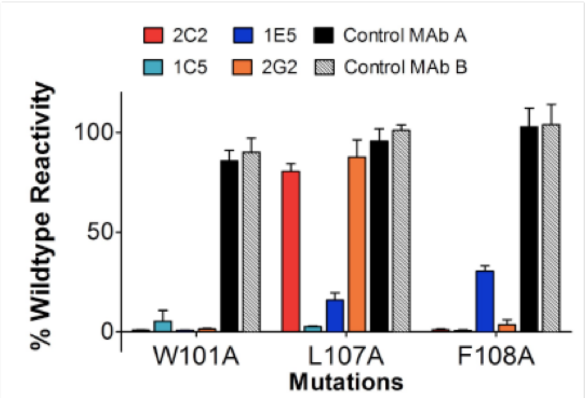


**Supplementary figure S1. Epitope mapping of the HuMAbs**

A) Identifying Critical Residues for MAbs 1C3, 1F11, 3E1, and 3G9. The DENV2 prM/E mutation library was assayed in an immunofluorescence flow cytometry assay, in duplicate, for binding by MAbs. Each raw data point was background-subtracted and normalized to the value for reactivity with wild-type DENV2 prM/E. We show graphs of 1C3 and 1F11 (representative of 1F11, 3E1, and 3G9) for the preliminary identification of critical clones. For each clone, the mean binding value is plotted as a function of the clone's mean expression value (gray circles), given by binding to a control MAb. We applied binding thresholds (dashed lines) for the test MAb (<20% of reactivity with wild-type prM/E) and a control MAb (>70% of reactivity with wild-type) to identify critical clones (red circles).

B) Identifying Critical Residues for MAbs 2C2, 2G2, 1C5, and 1E5. The HuMAbs were screened on DENV2 prM/E mutants with an FLE mutation, along with two control MAbs that do not bind the fusion loop region. The average binding values for each clone are shown as a percentage of binding to wild-type DEN2 prM/E. Clones with reactivity <20% relative to wild-type prM/E were identified as critical for MAb binding.

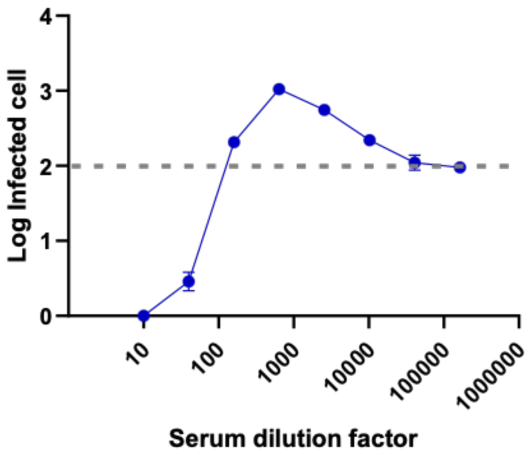


**Supplementary figure S2. ADE assay using DENV-2 NGC and DENV-2 immunized mouse serum**

Dotted line indicates the baseline of the infected cells in the control (100 infected cells; 2.0). Peak enhancement was observed at 1:640 diluted serum.
